# Lighting Up Mechanosensation: dyeing to see PIEZO2

**DOI:** 10.1101/2022.05.26.493600

**Authors:** Nicholas W. Villarino, Adrienne E. Dubin, Ardem Patapoutian, Kara L. Marshall

**Affiliations:** Department of Neuroscience, Dorris Neuroscience Center, The Scripps Research Institute, La Jolla, CA 92037; Howard Hughes Medical Institute, The Scripps Research Institute, La Jolla, CA 92037; Department of Neuroscience, Baylor College of Medicine, Houston TX; Jan and Dan Duncan Neurological Research Institute at Texas Children’s Hospital, Houston, TX, USA

**Author notes:** co-corresponding authors: A.P., K.L.M.

## Abstract

Sensory neurons gather information about mechanical forces from both the environment and internal organs to regulate physiology. PIEZO2 is a mechanosensory ion channel critical for touch, proprioception and bladder stretch sensation, yet its broad expression in sensory neurons and elsewhere suggests it may have undiscovered physiological roles. To fully understand mechanosensory physiology, we must know where and when PIEZO2-expressing neurons detect force. The fluorescent styryl dye FM 1-43 was shown to permeate several ion channels in cell culture and label sensory neurons. Surprisingly, we find that the vast majority of FM 1-43 somatosensory neuron labeling *in vivo* is dependent on PIEZO2-activity within nerve endings. We illustrate the potential of FM 1-43 by using it to identify novel *Piezo2*-expressing urethral neurons that are engaged by urination. These data reveal that FM 1-43 is a functional probe for mechanosensitivity via PIEZO2 activation *in vivo* and will facilitate the characterization of known and novel mechanosensory processes.

## RESULTS

Knowing when and where the nervous system detects mechanical force remains a challenge. The mechanosensory ion channel PIEZO2 is expressed in the peripheral nervous system, where it detects mechanical forces to mediate the senses of touch, proprioception, lung stretch, baroreception, and bladder fullness (Coste et al., 2010; Marshall et al., 2020; Ranade et al., 2014; Woo et al., 2015; Woo et al., 2014; Zeng et al., 2018). There are no known endogenous chemical agonists for PIEZO2, so identifying where PIEZO2 is expressed is an important clue that a specific cell or tissue may detect mechanical force. Unfortunately, antibodies against PIEZO2 or a PIEZO2-GFP knock-in mouse (Woo *et al*., 2015) have had limited efficacy for *in vivo* localization of PIEZO2. To aid in visualizing mechanosensory endings that might express PIEZO2, we turned to the fluorescent styryl dye FM 1-43 (Betz et al., 1996). This dye partitions into lipid membranes and has been used to study vesicular recycling, but also is proposed to enter cells through a separate mechanism: sensory ion channel permeation (Meyers et al., 2003). This feature has enabled the study of mechanotransduction ion channel activity in hair cells, and others have noted cell labeling in cell culture that is dependent on expression of TRPA1, TRPV1, and P2X ion channels (Karashima et al., 2010; Meyers *et al*., 2003). Following these discoveries, FM 1-43 was used as a nonspecific sensory cell labeling tool in a wide variety of contexts (Majumder et al., 2017; Maksimovic et al., 2014; Marshall et al., 2015; Moayedi et al., 2018). This dye is reported to permeate mechanosensory channels in sensory neurons of the dorsal root ganglia (Drew and Wood, 2007), indicating that it should label PIEZO2 positive endings.

We thus set out to determine how much of *in vivo* FM1-43 labeling in sensory neurons depends on PIEZO2, the primary mechanotransduction channel expressed by DRG neurons. Constitutive knockout of *Piezo2* in all tissues is lethal, so we used a Cre recombinase driven by the segmentation gene *HoxB8* to delete functional *Piezo2* from caudal tissues only (Witschi et al., 2010; Woo *et al*., 2015). *HoxB8*^*Cre*^ expression begins in the lower cervical regions and increases to full expression in thoracic levels (Witschi *et al*., 2010). Bright cell body labeling was seen in DRGs 24 hours after an intraperitoneal injection of FM1-43 (1.12 mg/kg). To our surprise, the widespread labeling of DRG cell bodies was almost completely absent from the caudal DRGs in *HoxB8*^*Cre+*^*;Piezo2*^*f/f*^ mice 24 h after FM1-43 injection (Fig. 1A). Some FM1-43 was visible in the vasculature (Supplemental Figure 1), but only a few cell bodies remained labeled in caudal *Piezo2* knockout DRGs. Some remaining labeling could be attributed to low residual *Piezo2* transcript observed in *HoxB8*^*Cre+*^*;Piezo2*^*f/f*^ mice (Murthy et al., 2018). We also used *Phox2b*^*Cre+*^*;Piezo2*^*f/f*^ animals to remove functional *Piezo2* from the nodose ganglion, which supplies sensory innervation to the major organs of the body via the vagus nerve. Nodose neuronal cell body labeling was virtually absent in knockout animals, whereas the labeling in the adjacent jugular ganglion, which is not targeted by *Phox2b*^*Cre*^, remained (Fig. 1B). Quantification of DRG labeling revealed a gradient of decreasing fluorescence in *HoxB8*^*Cre+*^*;Piezo2*^*f/f*^ knockout ganglia that corresponds well to the increasing rostral-to-caudal gradient of *HoxB8*^*Cre*^ expression (Fig. 1C). Together, these data reveal that the majority of FM1-43 sensory neuron labeling is dependent on PIEZO2 expression.

**Figure 1:**
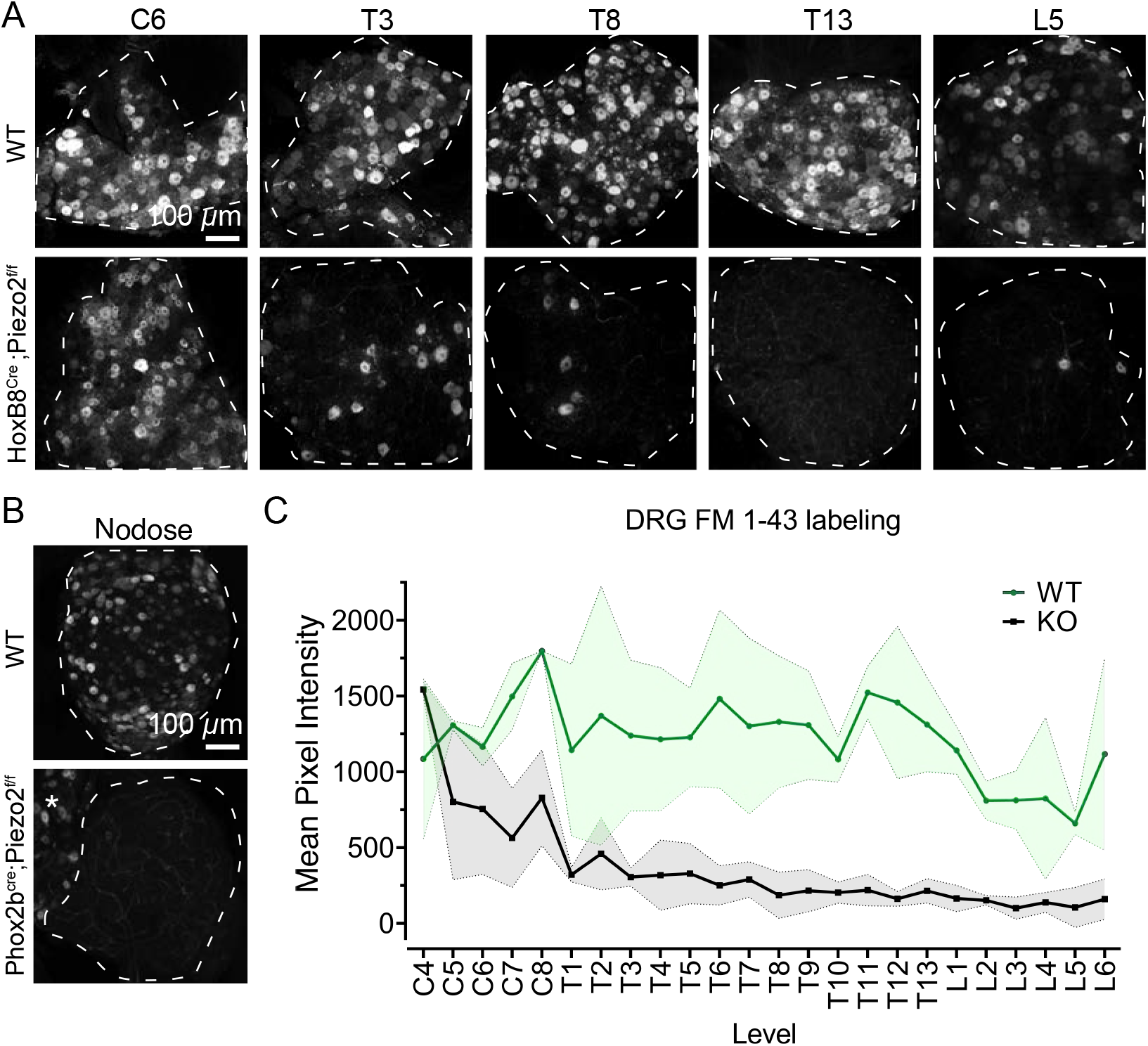
The majority of FM 1-43 sensory neuron uptake depends on *Piezo2* expression. (A) Representative z-stack images of whole-mount dorsal root ganglia DRG from wildtype (Top) or *HoxB8*^*Cre+*^*;Piezo2*^*f/f*^ (Bottom) mice that were injected i.p. with FM 1-43 24 h prior. The DRG level written on the top applies to both images underneath it. Scale bar applies to all images. Tissue area outlined with dashed line. (B) Representative z-stack images of whole-mount nodose ganglion from wildtype (Top) and *Phox2b*^*Cre+*^*;Piezo2*^*f/f*^ mice injected i.p. with FM 1-43 24 h prior. Asterisk indicates the jugular ganglion, which is fused to the nodose in mice. Nodose area outlined with dashed line. Scale bar applies to both images. (C) Quantification of mean fluorescence intensity of FM 1-43 labeling in DRGs at each vertebral levels from wildtype (N=2-3) and *HoxB8*^*Cre+*^*;Piezo2*^*f/f*^ KO (N=2-4) animals per level. Center line is mean, with shaded areas representing SD. Comparison of all lumbar values between WT and KO is significantly different. Unpaired two-tailed student’s t test, p < 0.0001

Mechanotransduction sites are located in sensory neuron endings, which are distant from cell bodies in the DRG and nodose ganglion. Injection of FM1-43 *in vivo* robustly labels sensory endings in addition to sensory neuron cell bodies. One example of robust FM 1-43 labeling occurs in lanceolate endings of the skin, which surround follicles to detect hair movement and convey the sense of touch (Abraira and Ginty, 2013) (Fig. 2A). Some circumferential endings and Merkel-cell neurite complexes are also labeled (Supplemental Figure 1). Consistent with what we observed in DRGs, lanceolate ending labeling on hind-leg skin was also absent in *HoxB8*^*Cre+*^*;Piezo2*^*f/f*^ mice (Fig. 2B). To investigate sensory endings supplied by the nodose ganglion, we surveyed the baroreceptor endings that wrap around the central arch of the aorta to detect blood pressure (Min et al., 2019; Zeng *et al*., 2018)(Fig. 2C). Min et al. and others have shown that PIEZO2 positive afferents are only a subset of neurons that innervate the aortic arch (Elsaafien et al., 2022). Consistent with this, we see a subset of aortic arch neurons labeled by FM1-43 (Supplemental Figure 1). These endings were unlabeled in *Phox2b*^*Cre+*^*;Piezo2*^*f/f*^ animals injected with FM1-43 (Fig. 2D). Our previous work showed that baroreceptor endings are still present in *Piezo2* knockout (KO) animals and still functional because of redundancy provided by the related mechanotransduction ion channel PIEZO1 (Zeng *et al*., 2018). Given that we see complete loss of aortic arch labeling and nodose neuron labeling in *Piezo2* KO animals, this suggests that *in vivo* FM 1-43 labeling in sensory neurons depends on *Piezo2* expression, but not *Piezo1*, which is expressed in some nodose neurons (Elsaafien *et al*., 2022).

**Figure 2:**
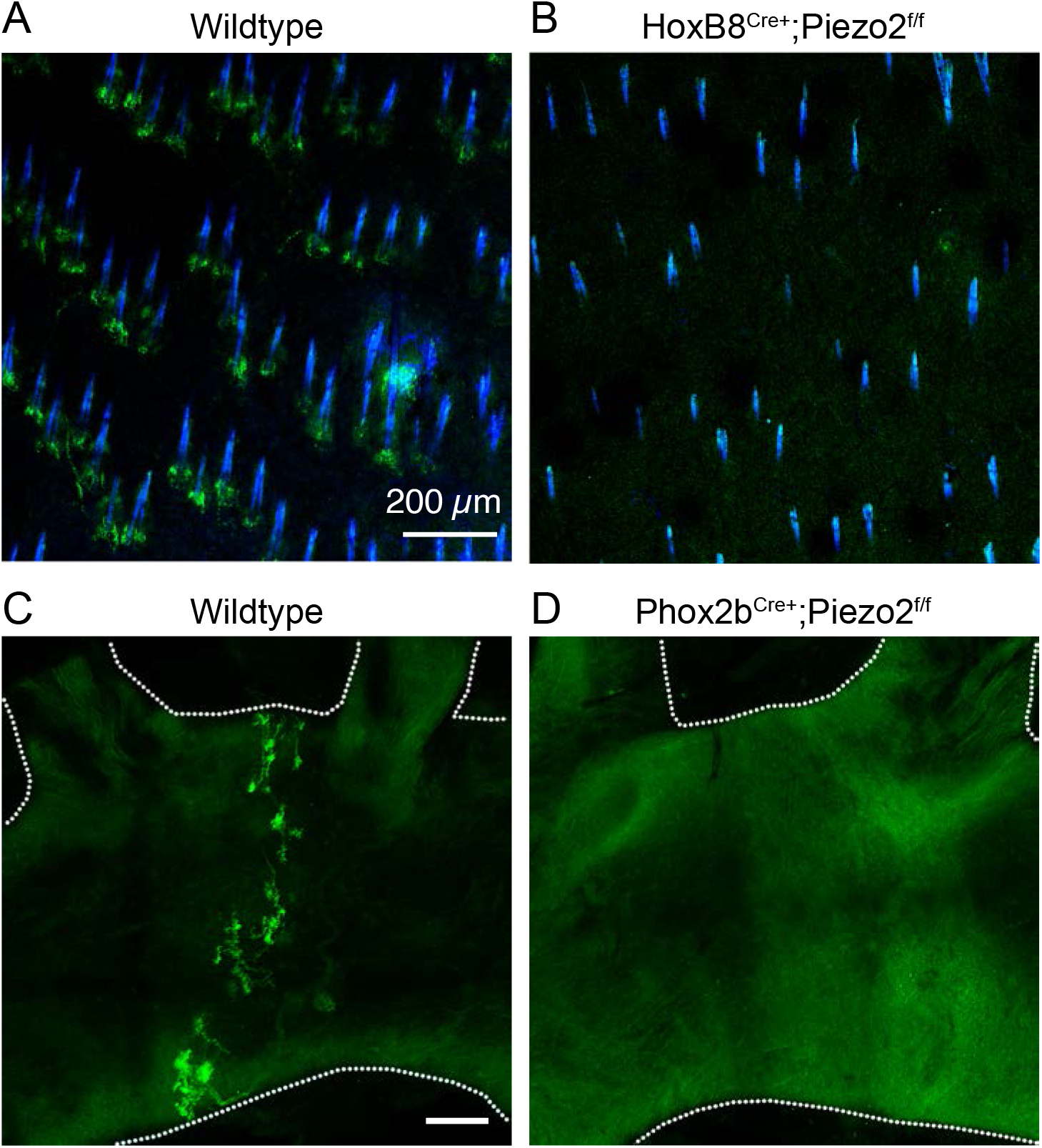
PIEZO2 is required for FM 1-43 labeling of lanceolate and aortic arch sensory endings. (A) Representative 24 h FM 1-43 labeling (green) of cutaneous neurons in whole-mount skin from wildtype animals. Scale applies to A and B. Hair shaft autofluorescence is shown in blue for A–B. (B) Representative 24 h FM 1-43 labeling of whole-mount skin from *HoxB8*^*Cre+*^*;Piezo2*^*f/f*^ KO mouse. (C) Representative 24 h FM 1-43 labeling of whole-mount aorta baroreceptor endings in wildtype animals. Scale bar: 200 μm, applies to C and D. Aorta tissue is outlined. (D) Representative 24 h FM 1-43 labeling of whole-mount aorta *Phox2b*^*Cre+*^*;Piezo2*^*f/f*^ knockout mouse.

If FM1-43 permeates PIEZO2-positive neurons at the site of mechanotransduction, then it should enter first at the sensory endings, where mechanical stimuli are detected. On a short timescale in an awake, behaving animal, sensory neuron endings of varied functions would be activated with different frequencies, or sometimes not at all. To understand if labeling can occur rapidly in active sensory endings, we visualized the baroreceptor endings at different time points. These endings are activated *in vivo* with each heartbeat (Zeng *et al*., 2018). Consistent with this ongoing activation, we observed FM1-43 labeling in the baroreceptor endings only 90 minutes after intraperitoneal injection. At this time point there were no cell bodies labeled in the nodose ganglion (Fig. 3A). The dye should only reach cell bodies after sufficient time has passed for trafficking back from the periphery. As predicted, baroreceptor endings were more brightly fluorescent after 6 h, and some nodose ganglion cell bodies were labeled (Fig. 3A). The aortic arch is in the chest cavity near the nodose ganglion cell bodies. For many sensory neurons, target tissues in the body are far from their vertebral or cranial ganglia. Evidence that cell body labeling is due to trafficking from endings is supported by our observation that FM1-43 accumulates over longer timescales in DRGs (Fig. 3B). Cell bodies were dimly labeled 12 h after injection, but once labeled they displayed remarkable persistence; bright labeling remained even 7 weeks after injection (Fig. 3B). Together these data indicate that FM 1-43 labeling is first occurring at sites of sensory transduction before being trafficked back to cell bodies, where labeling persists.

**Figure 3:**
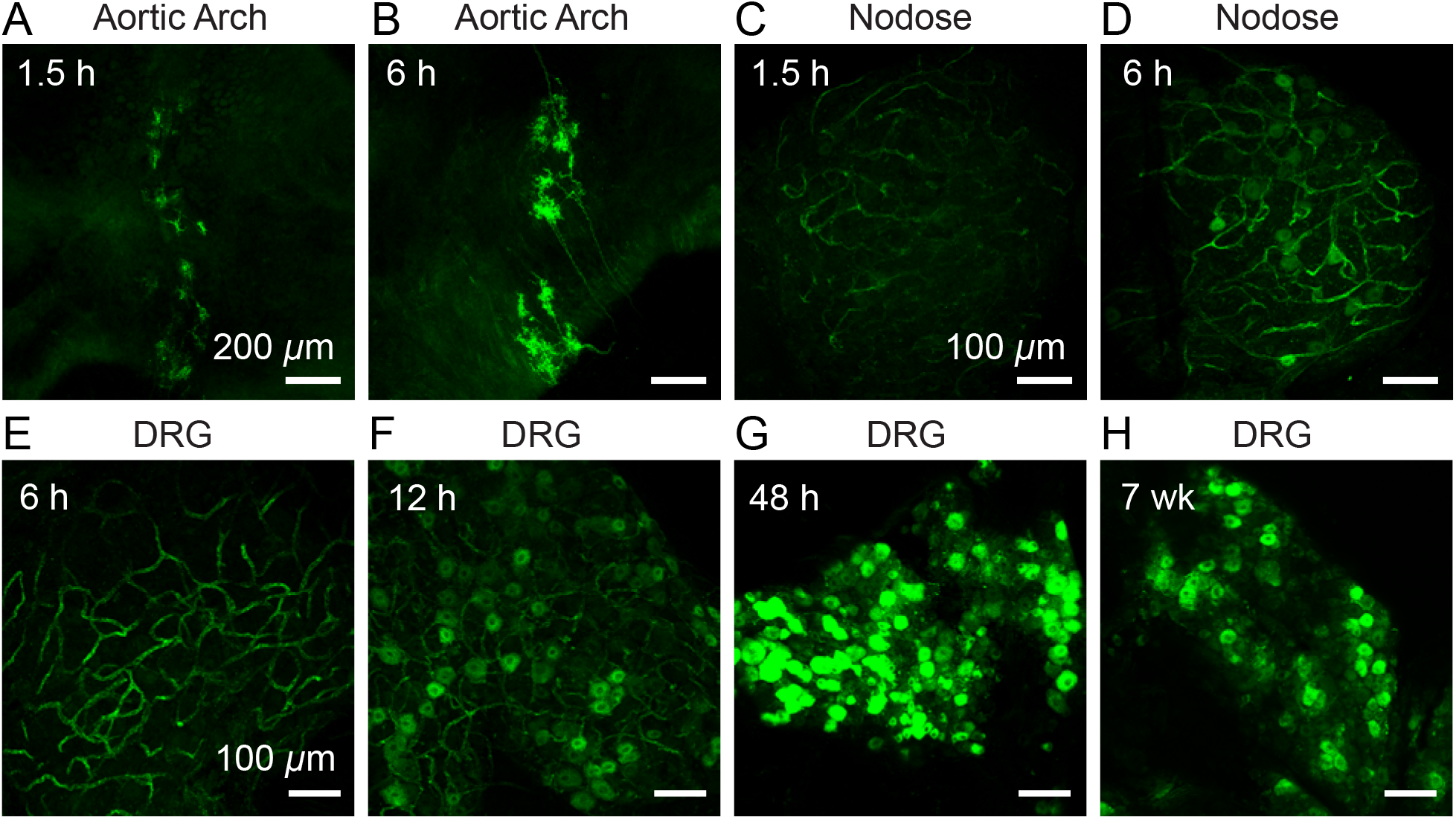
Labeling of sensory cell bodies occurs in a time-dependent manner and persists for weeks. (A) Representative FM 1-43 labeling in the aorta baroreceptor ending after 1.5 h of FM 1-43 labeling. Scale also applies to B. (B) Representative FM 1-43 labeling in the aorta baroreceptor ending after 6 h of FM 1-43 labeling. (C) Representative FM 1-43 labeling in the nodose ganglion from the same animal shown in A after 1.5 h of FM 1-43 labeling. Scale also applies to D. (D) Representative FM 1-43 labeling in the nodose ganglion from the same animal shown in B after 6 h of FM 1-43 labeling. (E) Representative z-stack of whole-mount lumbar DRG from wildtype mice injected i.p. with FM 1-43 6 h before tissue collection. Scale applies to E–H. (F) Representative z-stack of whole-mount lumbar DRG from wildtype mice injected i.p. with FM 1-43 12h before tissue collection. (G) Representative z-stack of whole-mount lumbar DRG from wildtype mice injected i.p. with FM 1-43 48 h before tissue collection. (H) Representative z-stack of whole-mount lumbar DRG from wildtype mice injected i.p. with FM 1-43 7 weeks before tissue collection.

Our observation that FM1-43 uptake initiates at the sites of sensory transduction led us to ask whether sensory neuron labeling requires PIEZO2-dependent activity. Gentle touch is almost entirely dependent on PIEZO2 (Chesler et al., 2016; Ranade *et al*., 2014), so we used this established role to test the *in vivo* activity dependence of FM1-43 in touch neurons. Lanceolate endings form fence-like structures surrounding the base of hair follicles, and they respond to hair deflection with exquisite sensitivity (Handler and Ginty, 2021). We reasoned that removal of external hair shafts on a patch of leg skin would reduce optimal PIEZO2-dependent mechanical activation of lanceolate afferents, and thus suppress FM 1-43 loading in this region. Remarkably, minimal labeling of lanceolate endings was present in de-haired hind-leg skin after 24 h, whereas the hairy regions in the contralateral control leg displayed bright labeling (Fig. 4A–C). These results suggest that *Piezo2-*dependent activity is necessary for robust dye labeling *in vivo*.

**Figure 4:**
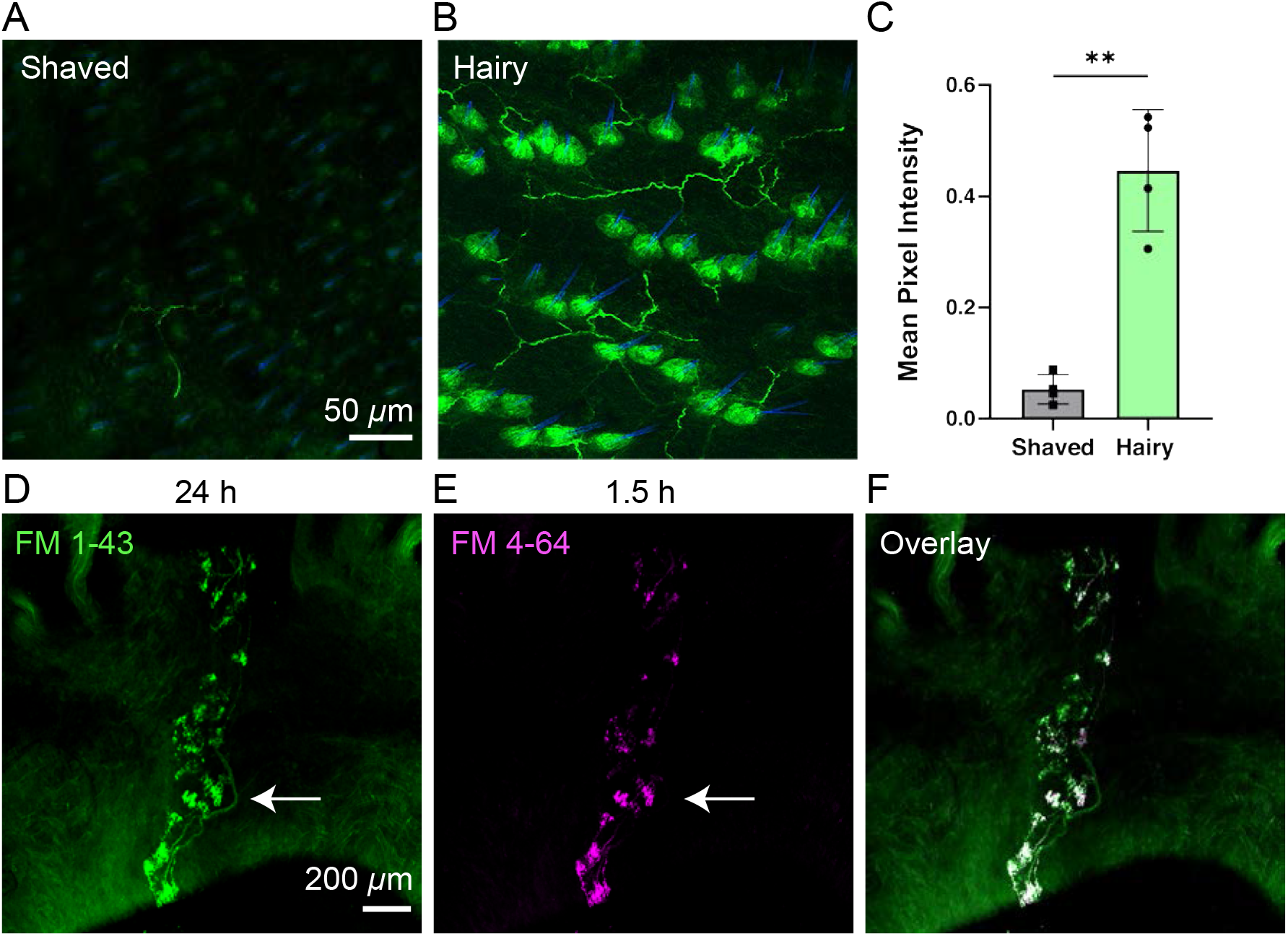
FM dyes capture activity patterns of complex sensory ending structures. (A) Representative fluorescence image from ‘Shaved’ de-haired skin. Scale bar applies to A and B. (B) Representative fluorescence image of ‘Hairy’ skin at 24 h after two i.p. FM 1-43 injections, 8 h apart. Lanceolate endings around the hair follicles are brightly labeled with additional halo of signal in the surrounding follicle. Follicle signal was only seen in skin after two injections of FM 1-43. (C) Quantification of mean fluorescence from FM 1-43 signal normalized to hair follicle count, which were visualized through blue autofluorescence of hair shafts. Individual dots represent mean values from 4-6 fields of view from each condition per mouse. N=4 mice were analyzed per group using a paired *t*-test (p = 0.0035). (D) FM 1-43 labeling in a whole-mount aorta from an animal injected with FM 1-43 24 h before tissue harvest. Arrow labels nerve trunk. Scale bar applies to D–F. (E) The same baroreceptor endings in D, labeled by FM 4-64 injected 1.5 h before tissue harvest. (F) Merge of D and E. White represents areas of overlap.

Activity dependence of FM 1-43 dye labeling indicates that it might be used in an animal to assess which neurons or specific neuron-ending structures are most engaged during a given time period. To test this, we injected mice with FM 1-43 24 h before harvesting to label the full neuron ending structure and nerve trunk, which only labels after sufficient time for trafficking and diffusion (Fig. 4D). In the same animal we injected the red-shifted FM analog, FM 4-64, 1.5 h before euthanizing (Fig. 4E). FM 1-43 and FM 4-64 have different fluorescent emission spectra, which allows distinguishing these dyes within the same cell. Critically, FM 4-64 labeling of sensory neurons is also *Piezo2-*dependent (Supplemental Figure 2). This approach revealed that the *Piezo2-*expressing endings in the aortic arch, shown by others to be most prominent near the arterial ligament (Min *et al*., 2019), labeled robustly at 90 min, while structures toward the top of the aortic arch were not as well labeled. As expected, the nerve trunk feeding these endings was only labeled from the 24 h injection (Fig. 4F). Thus, labeling the same neuron at multiple time points can identify precise regions where the most activity occurred. This technique offers a novel methodology for parsing the activity landscape of complex, interwoven endings.

To further investigate whether FM1-43 labeling depends on PIEZO2 activity, we tested dye uptake into cells heterologously expressing *PIEZO2* in cell culture. For all cell culture assays related to mechanosensitivity, we use HEK cells engineered to lack endogenous PIEZO1 (HEK P1KO) to avoid potential confounding effects (Dubin et al., 2017). Two days after transfection with a *PIEZO2* cDNA plasmid, HEK P1KO cells were treated with 10 μM FM 1-43 for 20 minutes followed by washing with Advasep-7, a β-cyclodextrin known to reduce background signal from FM 1-43 (Kay et al., 1999). Washed cells were dissociated and subjected to flow cytometry for quantification of population fluorescence changes (Fig. 5). FM 1-43 robustly labeled cells overexpressing PIEZO2 but did not label cells transfected with control plasmid (Fig. 5A–F). Additionally, we subjected cells to orbital shaking as a simple attempt to increase mechanical stimulation and observed that labeling increased by 49 ± 8%, suggesting stimulus-dependent uptake (Fig. 5J). Notably, even in the absence of mechanical stimulation, cells overexpressing PIEZO2 readily labeled. A similar result was observed with TRPV1 overexpression, indicating that some dye permeation still occurs even without canonical channel activating stimuli (Supplemental Figure 3). Importantly, fluorescence in control cells was unchanged with orbital shaking (Fig. 5A–C) ruling out the possibility that stimulation alone increased background labeling.

**Figure 5:**
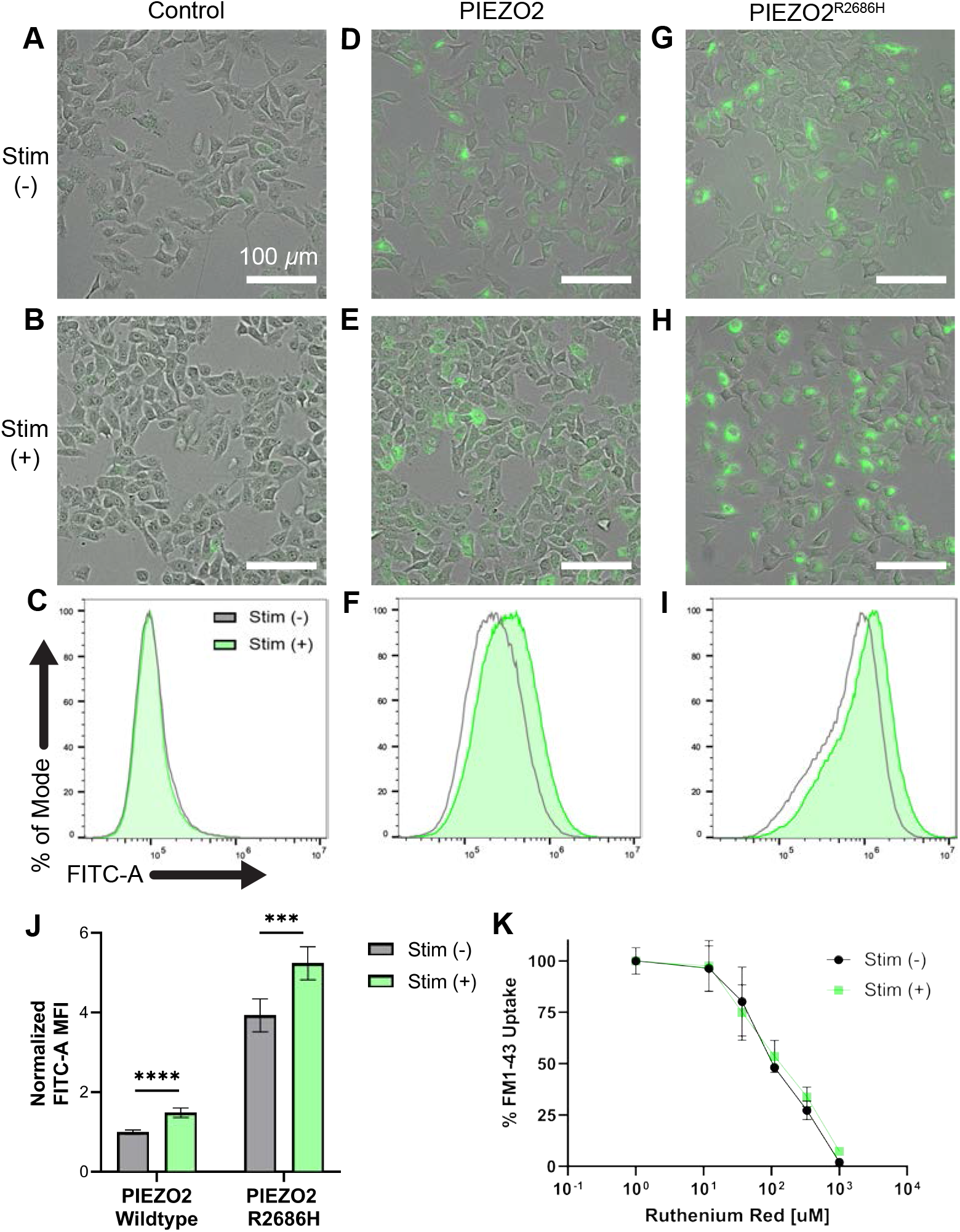
In cell culture, FM 1-43 labeling increases with human PIEZO2 channel activity. (A) Representative images from HEK P1KO cells that were transfected with control plasmid, exposed to FM 1-43, and had no stimulus. Scale applies to all images. (B) HEK P1KO cells transfected with control plasmid and exposed to orbital shaking. (C) Representative flow cytometry plots from HEK P1KO cells transfected with control plasmid. X-axis is FITC-A (log scale) and represents FM 1-43 loading; Y-axis is % cell count. (D) Images from HEK P1KO cells transfected with *PIEZO2* with no stimulus. (E) HEK P1KO cells transfected with *PIEZO2* and exposed to orbital shaking. (F) Flow cytometry histograms of *PIEZO2* transfected HEK P1KO cells with (green) and without (gray) exposure to shear stress. X-axis is FITC-A (log scale) and represents FM 1-43 loading; Y-axis is % cell count. (G) Representative images and flow cytometry plots from HEK cells transfected with *PIEZO2*^*R2868H*^, a gain-of-function allele, with no stimulus. (H) HEK P1KO cells transfected with the *PIEZO2*^*R2868H*^ and exposed to orbital shaking. (I) Representative flow cytometry histograms from HEK cells transfected with *PIEZO2*^*R2868H*^ with (green) and without (gray) exposure to orbital shaking. X-axis is FITC-A (log scale) and represents FM 1-43 loading; Y-axis is % cell count. (J) Flow cytometry quantification of each transfection condition with and without orbital shaking, normalized to control transfected cells. Means are averaged from 3 FACS experiments with 1-3 biological replicates for a total of 6 population averages per group. Unpaired Student’s *t*-test, PIEZO2 WT p-value = 0.00003 and PIEZO2^R2686H^ p-value = 0.0002. (K) FM 1-43 uptake, shown as a percent of maximum, is suppressed in a concentration-dependent manner during exposure to ruthenium red (x-axis).

To further assess the relationship between FM1-43 uptake and PIEZO2 channel activity, we transfected *PIEZO2*^*R2686H*^ gain-of-function cDNA. This mutation results in slowed current inactivation kinetics during poke-induced channel activation, increasing the time constant of inactivation by 435 ± 36% (Supplemental Figure 4). This gain of function mutation has been causally linked to Gordon syndrome and distal arthrogryposis type 5, two rare autosomal dominant multisystem disorders observed in human patients (Bagriantsev et al., 2014; Wu et al., 2017) Consistent with this, we observed a 393 ± 10% increase in fluorescence in *PIEZO2*^*R2686H*^ transfected cells over unstimulated *PIEZO2* WT cells, and mechanically stimulated *PIEZO2*^*R2686H*^ exhibited a 35 ± 10% increase in fluorescence compared to the unstimulated condition (Fig. 5G–J).

To confirm that FM1-43 entry is attributable to ion channel pore opening, we tested dye uptake in the presence the non-selective ion channel blocker Ruthenium Red (Malecot et al., 1998; Szallasi and Blumberg, 1999). Concentration-dependent block of fluorescence with and without mechanical stimulation was observed with IC_50_ values of 115 ± 27 μM and 135 ± 31 μM, respectively (Fig. 5K). We previously reported 30–50 μM was sufficient to block of PIEZO2-dependent mechanically activated currents in sensory neurons using acute electrophysiological recordings (Coste *et al*., 2010; Shin et al., 2019). It is possible the higher IC_50_ values were due to us monitoring FM 1-43 uptake in a field of cells where labeling represents cumulative PIEZO2 activity over a longer duration than is typically monitored in an electrophysiological setting. Together, these data reveal a direct relationship between PIEZO2 channel activity and dye uptake.

The activity dependence of FM1-43 labeling in awake, behaving mice presents a new opportunity to discover novel sensory neurons and physiological processes that rely on PIEZO2. Previously, we found that PIEZO2 was critical for the sensation of bladder fullness and setting urinary reflexes, and we aimed to use FM 1-43 to identify PIEZO2-positive neurons in the lower urinary tract (LUT) that could mediate these functions (Marshall *et al*., 2020). Labeled neuron-like endings were observed in the bladder muscle and more interior layers (Supplemental Figure 5) but were sometimes difficult to visualize in whole-mount against the high background, in part arising from urothelial labeling (Supplemental Figure 5). It is possible that urothelium labeling is due to the large amount of vesicular recycling that occurs during bladder stretch (Truschel et al., 2002). Bladder neurons were less evident in *HoxB8*^*Cre+*^*;Piezo2*^*f/f*^ knockout mice compared to wildtype mice (Supplemental Figure 5), but neuron labeling varied between wildtype animals and bladder region, making direct comparison difficult.

Neurons that could be involved in detecting bladder stretch were difficult to visualize, but mechanosensory neurons associated with the urethra also contribute to urinary reflexes and are not as well studied (de Groat et al., 2015). Our survey of the LUT with FM1-43 revealed a novel population of neurons in the urethra. We noticed labeling of dense neuronal endings along the entirety of the male bulbous urethra (Fig. 6A), the region between muscular attachments to the pelvic bone and the curve that begins the penile urethra. These endings were only found on the lateral aspects of the male urethral shaft (Fig. 6B–C). When viewed from the side, they have a distinctive tree-like morphology, and we have termed them “Torrey Pines” neurons due to their striking resemblance to the Torrey Pine, a species of pine tree uniquely native to San Diego and Santa Rosa Island (Fig 6D and Supplemental Fig. 6). These neurons failed to label in *HoxB8*^*Cre+;*^*Piezo2*^*f/f*^ mice (Fig. 6E), indicating that, as with other sensory neuron endings we investigated, their labeling depends on PIEZO2. Morphologically distinct neuron endings, which looked like simple free nerve endings, were also observed in the distal urethra of males and females (Fig. 6A, Supplemental Figure 6). Furthermore, using the dual-labeling approach described in Figure 4, we determined that the “leaves” of the trees are the sites of initial dye entry, and are thus plausible sites of transduction (Fig. 6F–H).

**Figure 6:**
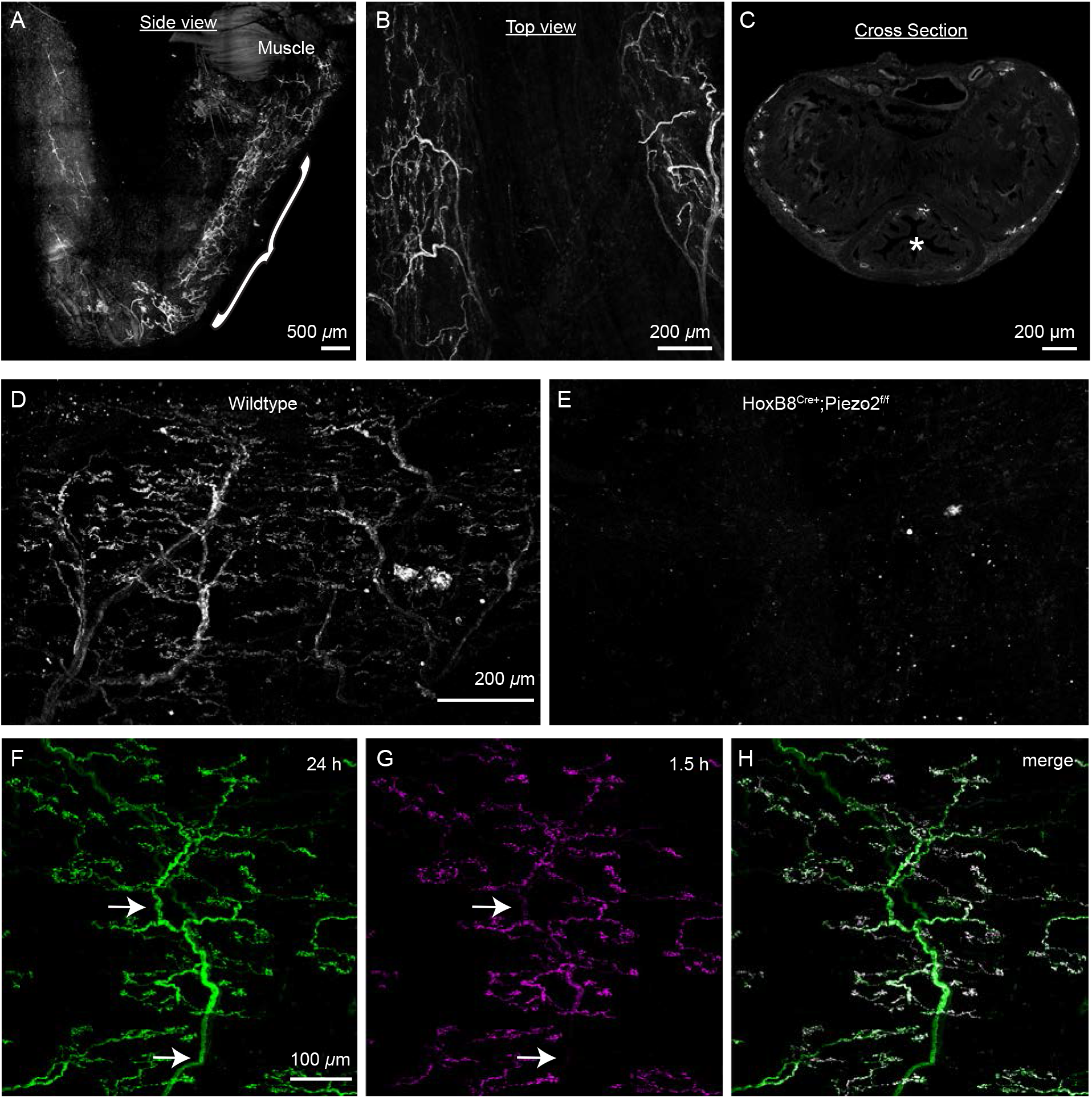
FM 1-43 reveals novel lateralized sensory endings in the bulbous urethra of male mice. (A) Side view of the male bulbous urethra labeled by FM 1-43 injection 24 h before tissue harvest. Bracket indicates bulbous urethra and area of “Torrey Pines” neuron labeling. Note free nerve endings in the most distal penile urethra. (B)Top view of the bulbous urethra (C)Cross-sectional view of the bulbous urethra shaft showing labeling around the edges. Urethral opening is marked by an asterisk. (D)24 h FM 1-43 labeling of “Torrey Pines” neurons in a wildtype bulbous urethra. (E)24 h after FM 1-43 injection in a *HoxB8*^*Cre+*^*;Piezo2*^*f/f*^ KO mouse, the bulbous urethra shows no neuronal labeling. (F)24 h labeling of a Torrey Pine ending shows nerve trunks (arrows). (G)1.5 h labeling of the same Torrey Pine neuron in F with FM 4-64 only shows labeling on the “leaves” of the tree. (H)Merged image from channels in F,G. White represents overlap.

In a preliminary effort to elucidate the molecular features of Torrey Pines neurons, we colocalized FM dye labeling with neuronal markers using whole-mount immunohistochemistry and genetic reporter mice. Despite dense peptidergic innervation of the urinary tract (Barry et al., 2018; Bertrand et al., 2020), Torrey pines neurons were not CGRP-positive (Supplemental Figure 6). We also found Torrey Pines neurons were beta-III tubulin- and neurofilament heavy (NFH)-positive, indicating that these are medium to large diameter neurons (Fig. 7A–D). While the nerve trunks were positive for these structural proteins, immunolabeling did not extend to the endings, so the complex ending morphology was exclusively resolved by the dye. This could explain why this morphology has not been previously described (Forrest et al., 2014) and is an advantage of using FM1-43 for neuron localization.

**Figure 7:**
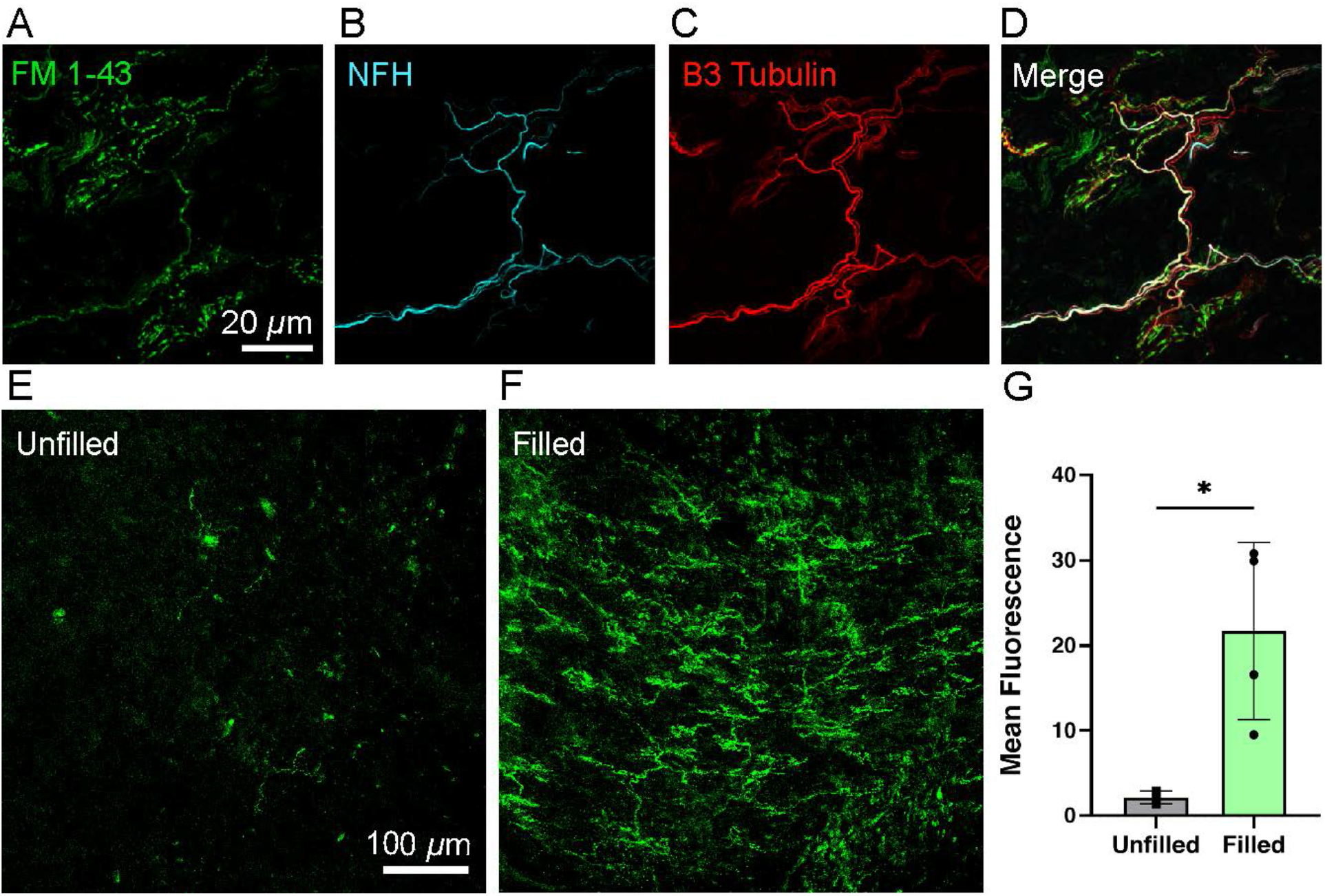
Characterization of Torrey Pines neurons, which are engaged during urination. (A) FM 1-43 labeling of a Torrey Pine neuron, 24 h after injection. (B) Neurofilament heavy (NFH) labeling of the Torrey Pine ending. (C) Beta-III tubulin labeling of the Torrey Pine ending. (D) Merge of channels in A–C. (E) Minimal FM 1-43 labeling is observed in the bulbous urethra when the bladder remains unfilled. (F) Robust Torrey pines neuron labeling is observed after 1 h of continuous bladder filling and recurring micturition cycles. (G) Quantification of FM 1-43 labeling brightness in filled versus unfilled animals. N=4 mice were analyzed per condition using an unpaired t-test (p < 0.024).

We next set out to identify the appropriate stimulus sufficient to induce FM 1-43 labeling in Torrey Pines neuronal endings in the urethra. We have previously shown that PIEZO2 is important for micturition reflexes and given their location, we hypothesized that these neurons would be active during urination. We injected FM1-43 i.p. in lightly anesthetized mice with a catheter inserted in the bladder as previously described, which allowed us to control fluid filling of the bladder (Marshall *et al*., 2020). When we continuously filled the bladder with saline, mice experienced repeated micturition events throughout the duration of the experiment, while control sham mice with unfilled bladders did not. Remarkably, Torrey Pines neuron labeling was prominent after 1 h of filling, and nearly absent in the unfilled condition (Fig. 7E–F). This identifies urination as a relevant stimulus that engages Torrey Pines neurons sufficiently to cause PIEZO2-dependent FM1-43 labeling.

## DISCUSSION

These results indicate that, given the selectivity of FM 1-43, it can be employed as an indicator of PIEZO2 activity in murine somatosensory endings and can be used to identify novel PIEZO2-dependent mechanosensitive processes. The apparent *in vivo* selectivity of FM 1-43 for PIEZO2 was a surprising finding, given that it has been proposed to permeate various ion channels in cell culture (Meyers *et al*., 2003). We believe that there are two possible explanations for this result. First, it is possible that that FM 1-43 is specific to PIEZO2 activity *in vivo* because other ion channels that are reported to facilitate FM1-43 permeation in cell culture, such as TRPV1 and P2X, are most active during pain or inflammatory conditions (Giniatullin, 2020; Karashima *et al*., 2010). Therefore, the vast majority of FM1-43 uptake is PIEZO2-dependent because this mechanosensory ion channel is the primary sensory ion channel engaged regularly during a healthy animal’s day. Consistent with this, we see minimal overlap of genetically labeled CGRP expressing cells with FM dyes in DRGs (Supplemental Figure 6). Some CGRP expressing DRG neurons express PIEZO2 (Murthy *et al*., 2018), but these are thought to primarily represent nociceptive populations and we would expect these to be most active during injury or inflammation (Cowie et al., 2018). However, at present our understanding of the activation dynamics of PIEZO2 in such populations is incomplete and requires further study. Another possibility is that FM1-43 uptake via these previously identified ion channels occurs more readily in culture systems than *in vivo*. This difference could arise if FM1-43 labeling via certain ion channels requires high protein expression, which is achieved in cell culture with protein overexpression.

An additional surprise regarding selectivity of FM1-43 was that we observed preferential uptake of FM1-43 via PIEZO2 compared to PIEZO1 *in vivo*, as evidenced by a lack of baroreceptor neuron labeling in *Phox2b*^*Cre*^;*Piezo2* knockout mice (Elsaafien *et al*., 2022; Min *et al*., 2019). Future structure-function studies of these related ion channels could explore potential mechanisms of FM1-43 uptake (Coste *et al*., 2010).

Despite the body of work indicating that FM1-43 can label cells expressing various sensory receptors, we still do not know if FM1-43 directly permeates these ion channels. FM1-43 has traditionally been used to visualize vesicular recycling associated with synaptic activity because it inserts into the lipid membrane. For example, neuronal uptake in motor neurons has been shown to occur through endocytosis (Betz et al., 1992; Gaffield et al., 2011). However, Myers et al. and others found that FM1-43 uptake in sensory cells did not depend on endocytosis and noted that sensory neuron FM1-43 labeling was brighter and more persistent than that of motor neuron uptake, suggesting that it occurs through a distinct mechanism (Li et al., 2009; Meyers et al., 2003). Furthermore, it has also been proposed that FM1-43 is capable of permeating mechanosensory ion channels in DRG neurons (Drew and Wood, 2007). Future work is required to determine whether PIEZO2-dependent cell labeling is through direct permeation of PIEZO2, or whether labeling occurs downstream of PIEZO2-dependent activity. Regardless, our work demonstrates the capability of using FM1-43 as a marker of PIEZO2-dependent activity. However, given other potential mechanisms for cellular uptake, FM1-43 labeling cannot serve as proof of *Piezo2* expression, particularly in cell types that experience frequent endocytic and exocytic activity, such as umbrella cells of the urothelium. This should be a consideration when using FM 1-43 as an *in vivo* labeling tool.

Our finding that the majority of *in vivo* sensory neuron FM1-43 labeling is dependent on PIEZO2 largely corroborates previous work. For example, Merkel cells label with FM 1-43 injection and express PIEZO2 (Maksimovic *et al*., 2014; Woo *et al*., 2014). Interestingly, Moayedi et al. noted that Piezo2-GFP labeling in the oral cavity corresponded to FM 1-43 labeling (Moayedi *et al*., 2018). Given the many sensory structures that have been shown to label with FM 1-43, we believe that our data can provide additional context to previous work, as labeling could suggest PIEZO2 function. For example, structures observed in bat wings using FM 1-43 might represent PIEZO2-dependent touch receptors (Marshall *et al*., 2015).

The finding of robust FM 1-43 labeling in species like bats and star-nosed moles highlights an advantage of this technique, as it can be used without specialized genetic tools (Marasco et al., 2006; Marshall *et al*., 2015). FM 1-43 can also be used to investigate organs where mechanosensory functions are still poorly understood. In all these cases, confirmation of *Piezo2*-dependence would still be required, but if confirmed, the dye offers a method to verify the spatial and temporal elements of PIEZO2 activity. To our knowledge, this is the only example of a tool that allows for permanent localization of the activity of a specific ion channel. Similarly, our data reveal robust FM1-43 uptake in cell cultures expressing PIEZO2. Taken together we expect that these dyes will aid researchers in confirming and better defining the many functions of PIEZO2.

As proof of its capability, we use FM 1-43 to pinpoint a unique neuron morphology in the urethra, that of the Torrey Pines neurons, which appear to be engaged during urination. Further work is needed to fully characterize the origin of these neurons and define their function in urinary tract physiology, but this finding represents an example of new directions and hypotheses that are enabled by *in vivo* FM 1-43 labeling.

## ACKNOWLEDGEMENTS

The authors thank the Patapoutian lab for helpful discussions, E. Lumpkin for early discussions, R. Hill, M. Odem, L. Drew and A. Chesler for comments on the manuscript, and K. Spencer in the Nikon Center of Excellence at Scripps Research for microscopy help. This work was supported by the Howard Hughes Medical Institute; the NIH grants R35 NS105067 to A.P., F32 DK121494 and K99 DK128621 to K.L.M.

## METHODS

### Cloning

The full-length sequence of PIEZO2 transcript variant 2 (10520bp) was codon-optimized using the Thermo Fisher Gene Art Codon Optimization tool. G-blocks for this sequence were then generated by Azenta Life Sciences and were assembled using Gibson Assemble master mix (NEB: E2611) into a mammalian expression vector modified from pCDNA3.1 to contain a T2A self-cleaving peptide followed by an in-frame blue fluorescent protein (BFP) and a woodchuck hepatitis virus post-transcriptional regulatory element (WPRE). The final construct yielded was pCDNA3.1-*PIEZO2*-T2A-BFP-WPRE (“*PIEZO2”)*.

To generate the human *PIEZO2*-R2686H gain-of-function construct, the above sequence was modified using Q5 site-directed mutagenesis (E0554) directed at amino acid 2686 to substitute histidine for arginine, yielding pCDNA3.1-*PIEZO2_R2686H*-T2A-BFP-WPRE (“*PIEZO2*^*R2686H*^)

### Cell culture and transfection

The HEK P1KO cell line was maintained in DMEM (4.5 mg/ml glucose) with 10% FBS and pen/strep antibiotics. Prior to transfection, cells were plated on Matrigel (Corning: 354234), 75-90 μg/mL, coated Falcon Polystyrene 6-well tissue culture plates. When cells reached 50-70% confluency, cDNA was transfected into cells using PEI MAX at 1 μg/μL. 2 μg of DNA was transfected per well employing a ratio of 3:1 PEI (μg) to DNA (μg). Cells were transfected two days prior to FM1-43 labeling experiments. Cells from control condition were transfected with an empty vector containing BFP.

### FM1-43 Cell Culture Assay

For cell culture labeling experiments, FM1-43FX (Invitrogen: 35355) was resuspended at 10 μM in Hanks Buffered Saline Solution (HBSS, 10 mM HEPES, pH7.4). ADVASEP-7 (Biotium) was also resuspended at 500 μM in HBSS. Two days after transfection, cells were washed with HBSS and 10 μM FM1-43 was applied to cells. For stimulated condition, cells were placed on an orbital shaker (setting 6-7 out of 10), while unstimulated cells were left on a benchtop, both at room temperature. After 20 minutes, cells were washed twice with 500 μM ADVASEP-7, 3 minutes per wash with orbital shaking.

For FM1-43 labeling block experiments, Ruthenium Red (RR, Sigma: R2751) was resuspended in HBSS at stated concentrations (Fig 4). Cells were preincubated with RR for 10 min and contained RR at the indicated concentrations during FM1-43 labeling. In addition to two ADVASEP-7 washes, cells were washed twice more with HBSS to remove residual RR. To generate Ruthenium Red IC50 values in Prism (Graphpad), a nonlinear regression model using the ‘Dose-response – Inhibition’ ([Inhibitor] vs. normalized response) equation was fit to the data assuming a standard slope (Hill Slope = -1.0).

### Electrophysiology

Mechanically activated currents from HEK P1KO cells 2 days after transfection were recorded in whole-cell patch clamp mode using a MultiClamp700A amplifier and DigiData1550 (Molecular Devices) and stored directly and digitized online using pClamp software (version 10.7). Currents were sampled at 20 kHz and filtered at 2 kHz. Recording electrodes had a resistance of 2 to 5 megohms when filled with CsCl-based intracellular solution: 133 mM CsCl, 1 mM CaCl_2_, 1 mM MgCl_2_, 10 mM HEPES (pH with CsOH), 5 mM EGTA, 4 mM Mg-ATP (adenosine triphosphate), and 0.5 Na-GTP (guanosine triphosphate). Extracellular bath solution was composed of 133 mM NaCl, 3 mM KCl, 2.5 mM CaCl_2_, 1 mM MgCl_2_, 10 mM HEPES (pH 7.3 with NaOH), and 10 mM glucose. Mechanical stimulation was achieved using a fire-polished glass pipette (tip diameter, 3 to 4 μm) positioned at an angle of 80° relative to the cell being recorded. Displacement of the probe toward the cell was driven by Clampex-controlled piezoelectric crystal microstage (E625 LVPZT Controller/Amplifier; Physik Instrumente). The probe had a velocity of 1 μm ms−1 during the ramp phase of the command for forward movement, and the stimulus was applied for a duration of 125 ms. For each cell, a series of mechanical steps in 1 μm increments was applied every 10 s.

### Fluorescence activated cell sorting and quantification

For quantitative fluorescence analysis of FM 1-43 labeled HEK cells, cells were dissociated with 0.05% Trypsin-EDTA (Gibco: 25300054), washed once in FACS buffer (PBS supplemented with 2% FBS and 1 mM EDTA), resuspended in FACS buffer at 100K cells per mL, and strained into a 5 mL polystyrene Round-Bottom tube (Corning: 352235). Samples were then analyzed using the ACEA NovoCyte 3000 with NovoSampler Pro. Cells transfected with *PIEZO2* and *PIEZO2-R2686H* cDNA were gated on BFP, and subsequently FITC-A population mean fluorescent intensities were used as a fluorescent readout of FM 1-43 labeling. Data were analyzed using FlowJo (BD). Mean fluorescence values from ‘Control’ were interpreted as background labeling and were subtracted from mean fluorescence values of the experimental conditions. Remaining values were then normalized to the ‘PIEZO2 WT Unstimulated’ condition which operated as an internal control for comparing results between separate experiments. Three FACS experiments were run per group with 1-3 biological replicates per experiment, for a total of 6 population averages that were compared per condition.

### Mice

Mice were kept in standard housing with 12 h light/dark cycle set with lights on from 6 AM to 6 PM, with room temperature kept around 72 degrees Fahrenheit, with humidity between 30-80 % (not controlled). All mice used were adult male or female mice, as indicated in the text. The *HoxB8*^*Cre*^*;Piezo*^*f/f*^ and *Phox2b*^*Cre*^*;Piezo2*^*f/f*^ mouse lines have been previously described (Woo *et al*., 2015; Zeng *et al*., 2018) Genotyping was performed using guidelines from Jackson Laboratory. C57BL/6J adult mice were used for all experiments if not otherwise defined in the text.

### Styryl fluorescent dye labeling in mice and tissue collection

Unless otherwise stated, mice were injected intraperitoneally with FM1-43FX (Invitrogen: F35355) or FM 4-64FX (Invitrogen: F34653), at a dose of 1.12 mg/kg of body weight from stocks made up in PBS. At time post-injection time point indicated in the text, mice were perfused with PBS followed by 4% PFA. Tissues were subsequently dissected and drop fixed for a minimum of 1 h (smaller tissues) at 4°C and up to 24 h (larger tissues) at 4°C prior to imaging or immunostaining.

For urethra sectioning, tissue was fixed in 4% PFA for 24 h, embedded in Tissue Tek OCT (Sakura: 4583), and frozen. Using the cryostat, 30 μm cross sections throughout the bulbous urethra were generated onto Superfrost Plus microscope slides (Fisher: 12-550-15). Sections were then washed with PBS to remove OCT before mounting or staining.

### Cystometry

C57BL/6J male mice were anesthetized by isoflurane (3% induction, 1.1-1.4% maintenance, Kent Scientific SomnoSuite) and the bladder was catheterized and connected to saline lines, as described previously (Marshall *et al*., 2020). Saline lines were connected to a pressure sensor (Biopac: RX104A-MRI) which connected via pressure transducer (TSD104A) to an MP160 Biopac system amplifier (DA100C). Using a syringe pump, the bladder was continuously filled at 30 μl min^−1^ to induce urination events in the ‘filled’ condition. Mice in ‘unfilled’ condition were prepared in the same manner but did not receive continuous saline bladder filling. Filling continued for 60 min, after which mice were euthanized, and perfused with PBS and 4% PFA. The urethra was then collected, drop fixed in 4% PFA for 1 h, and imaged in a whole mount preparation.

### Whole mount immunohistochemistry

Bulbous urethra whole-mount immunostaining was performed as previously described for skin (Marshall et al., 2016). Tissues were collected from 8-12 week-old wildtype C57BL/6J mice after perfusion with 4% PFA. Tissues were fixed overnight in 4% PFA, washed 3 times in PBS, and washed for 5 h in approximately 40 mL of PBS with 0.1% TritonX-100 (PBSt), changing the PBSt every hour. Tissues were then placed in blocking solution consisting of 5% normal goat serum (NGS), 20% DMSO, 75% PBSt, along with primary antibodies. The primary antibodies used were chicken anti-neurofilament heavy chain (NFH) (1:500 dilution, Abcam: ab4680) and rabbit anti-beta III tubulin (1:500 dilution, Abcam: ab18207). Tissue was incubated for 4 days at 4°C on a rocker. Next the samples were washed in a large volume of PBSt for 5 h, changing PBSt every hour. Samples were then placed in blocking solution containing secondary antibodies. The secondary antibodies used were goat anti-chicken AlexaFluor 647 (Invitrogen A21449) and donkey anti-rabbit AlexaFluor 594 (A21207), all diluted at 1:500. Tissues were incubated for 2 days at 4°C on a rocker, with final washes in a PBSt for at least 5 h, with frequent PBSt changes. Samples were then blotted dry on a clean Kimwipe (Kimtech) and were mounted onto Superfrost Plus microscope slides.

### Activity dependent skin labeling

For the comparison of de-haired versus hairy skin, mice were lightly anesthetized, and just one hindlimb was shaved with clippers. This was followed by a brief exposure to Nair depilatory cream to completely remove external hair shafts. After 24 h to allow time for any skin sensitivity from depilation to recover, mice were injected i.p. with FM 1-43 as detailed above. To increase labeling, a second injection was given approximately 16 h later. Tissue was collected 24 h after initial injection, as described above. Hairy leg skin was first shaved and exposed to Nair before all skin was removed, washed in PBS and drop-fixed in 4% PFA. A scalpel was used to gently scrape the dermal fat away from skin to decrease background fluorescence before whole skin was mounted on slides for imaging.

### Confocal imaging

Prior to imaging, tissue was covered in Fluoromount G mounting medium with DAPI (Thermo: 00-4959-52) and number 1.5 coverglass was placed over the tissue. All confocal settings for a given experiment were imaged on the same microscope with identical settings for direct comparison of fluorescent brightness. Tissues were imaged on a Nikon C2 confocal mounted on an upright 90i microscope or an inverted NikonA1 confocal. For the Nikon C2, the following objectives were used: 10x PlanApo 0.45 Dry, 20X PlanApo, 0.8 Dry, 40X PlanFluor 1.3 Oil, and 60x PlanApo VC1.4 Oil. Laser lines used include 408nm, 488nm, 561nm and 637nm. Nikon C2 emission filters include 450/40nm, 514/30nm, and 585/65nm. For the NikonA1, the following objectives were used: 10x PlanApo 0.45. Laser lines included 405nm, 488nm, 561nm, and 638nm. Nikon A1 emission filters include 450/25nm, 525/25nm, 600/25nm, and 685/35nm.

### Image quantification

Images were analyzed in FIJI (ImageJ). For images of whole-mount DRGs, a region of interest was drawn around ganglion tissue, and mean brightness was measured. For Figure 1, An average background pixel brightness value was measured by taking mean intensity from an area without tissue from 36 images from both genotypes. This background value was subtracted from all brightness values. Tissue filled the area for some images, which is why an average background value was used. Mean brightness values at each vertebral level were quantified from 2-4 animals per level. C4-C6 is the range with only 2 animals per group. In most cases level brightness per animal used both left and right DRG images to calculate an average per animal, but when tissue was lost, there was only one image quantified per animal at that level.

For skin quantification, 4-6 images were taken around the hairy or shaved skin from each animal, and an average was calculated per animal. For quantification of urethra images, 3-5 images were taken evenly across the bulbous urethra region from each mouse to ensure a large portion of this region was captured even when no neuronal structures were visible. A threshold value was chosen from areas without labeling in order to reduce autofluorescence, and the same threshold was applied to all images in a given experiment. The entire image was filled by tissue, and the thresholded mean brightness of the entire image was measured. For skin images, mean brightness was normalized to the number of hair follicles in an image. The mean of all images was then taken for each animal, and these animal averages were compared between groups.

### Statistics

Graphing and statistical analysis was performed in Prism (GraphPad). Groups were compared using an unpaired Student’s two-tailed *t*-test, and a paired Student’s two-tailed *t*-test for Figure 4.

## Notes

### Competing Interest Statement

The authors have declared no competing interest.

## REFERENCES

Abraira, V.E., and Ginty, D.D. (2013). The sensory neurons of touch. Neuron 79, 618–639. 10.1016/j.neuron.2013.07.051.

Bagriantsev, S.N., Gracheva, E.O., and Gallagher, P.G. (2014). Piezo proteins: regulators of mechanosensation and other cellular processes. J Biol Chem 289, 31673–31681. 10.1074/jbc.R114.612697.

Barry, C.M., Ji, E., Sharma, H., Yap, P., Spencer, N.J., Matusica, D., and Haberberger, R.V. (2018). Peptidergic nerve fibers in the urethra: Morphological and neurochemical characteristics in female mice of reproductive age. Neurourol Urodyn 37, 960–970. 10.1002/nau.23434.

Bertrand, M.M., Korajkic, N., Osborne, P.B., and Keast, J.R. (2020). Functional segregation within the pelvic nerve of male rats: a meso- and microscopic analysis. J Anat 237, 757–773. 10.1111/joa.13221.

Betz, W.J., Mao, F., and Smith, C.B. (1996). Imaging exocytosis and endocytosis. Curr Opin Neurobiol 6, 365-371. Doi 10.1016/S0959-4388(96)80121-8.

Chesler, A.T., Szczot, M., Bharucha-Goebel, D., Ceko, M., Donkervoort, S., Laubacher, C., Hayes, L.H., Alter, K., Zampieri, C., Stanley, C., et al. (2016). The Role of PIEZO2 in Human Mechanosensation. N Engl J Med 375, 1355–1364. 10.1056/NEJMoa1602812.

Coste, B., Mathur, J., Schmidt, M., Earley, T.J., Ranade, S., Petrus, M.J., Dubin, A.E., and Patapoutian, A. (2010). Piezo1 and Piezo2 are essential components of distinct mechanically activated cation channels. Science 330, 55–60. 10.1126/science.1193270.

Cowie, A.M., Moehring, F., O’Hara, C., and Stucky, C.L. (2018). Optogenetic Inhibition of CGRPalpha Sensory Neurons Reveals Their Distinct Roles in Neuropathic and Incisional Pain. J Neurosci 38, 5807–5825. 10.1523/JNEUROSCI.3565-17.2018.

de Groat, W.C., Griffiths, D., and Yoshimura, N. (2015). Neural control of the lower urinary tract. Compr Physiol 5, 327–396. 10.1002/cphy.c130056.

Drew, L.J., and Wood, J.N. (2007). FM1-43 is a permeant blocker of mechanosensitive ion channels in sensory neurons and inhibits behavioural responses to mechanical stimuli. Mol Pain 3, 1. 10.1186/1744-8069-3-1.

Dubin, A.E., Murthy, S., Lewis, A.H., Brosse, L., Cahalan, S.M., Grandl, J., Coste, B., and Patapoutian, A. (2017). Endogenous Piezo1 Can Confound Mechanically Activated Channel Identification and Characterization. Neuron 94, 266–270 e263. 10.1016/j.neuron.2017.03.039.

Elsaafien, K., Harden, S.W., Johnson, D.N., Kimball, A.K., Sheng, W., Smith, J.A., Scott, K.A., Frazier, C.J., de Kloet, A.D., and Krause, E.G. (2022). A Novel Organ-Specific Approach to Selectively Target Sensory Afferents Innervating the Aortic Arch. Front Physiol 13, 841078. 10.3389/fphys.2022.841078.

Forrest, S.L., Osborne, P.B., and Keast, J.R. (2014). Characterization of axons expressing the artemin receptor in the female rat urinary bladder: a comparison with other major neuronal populations. J Comp Neurol 522, 3900–3927. 10.1002/cne.23648.

Giniatullin, R. (2020). Ion Channels of Nociception. Int J Mol Sci 21. 10.3390/ijms21103553.

Handler, A., and Ginty, D.D. (2021). The mechanosensory neurons of touch and their mechanisms of activation. Nat Rev Neurosci 22, 521–537. 10.1038/s41583-021-00489-x.

Karashima, Y., Prenen, J., Talavera, K., Janssens, A., Voets, T., and Nilius, B. (2010). Agonist-induced changes in Ca(2+) permeation through the nociceptor cation channel TRPA1. Biophys J 98, 773–783. 10.1016/j.bpj.2009.11.007.

Kay, A.R., Alfonso, A., Alford, S., Cline, H.T., Holgado, A.M., Sakmann, B., Snitsarev, V.A., Stricker, T.P., Takahashi, M., and Wu, L.G. (1999). Imaging synaptic activity in intact brain and slices with FM1-43 in C. elegans, lamprey, and rat. Neuron 24, 809–817. 10.1016/s0896-6273(00)81029-6.

Majumder, P., Moore, P.A., Richardson, G.P., and Gale, J.E. (2017). Protecting Mammalian Hair Cells from Aminoglycoside-Toxicity: Assessing Phenoxybenzamine’s Potential. Front Cell Neurosci 11, 94. 10.3389/fncel.2017.00094.

Maksimovic, S., Nakatani, M., Baba, Y., Nelson, A.M., Marshall, K.L., Wellnitz, S.A., Firozi, P., Woo, S.H., Ranade, S., Patapoutian, A., and Lumpkin, E.A. (2014). Epidermal Merkel cells are mechanosensory cells that tune mammalian touch receptors. Nature 509, 617–621. 10.1038/nature13250.

Malecot, C.O., Bito, V., and Argibay, J.A. (1998). Ruthenium red as an effective blocker of calcium and sodium currents in guinea-pig isolated ventricular heart cells. Br J Pharmacol 124, 465–472. 10.1038/sj.bjp.0701854.

Marasco, P.D., Tsuruda, P.R., Bautista, D.M., Julius, D., and Catania, K.C. (2006). Neuroanatomical evidence for segregation of nerve fibers conveying light touch and pain sensation in Eimer’s organ of the mole. Proc Natl Acad Sci U S A 103, 9339–9344. 10.1073/pnas.0603229103.

Marshall, K.L., Chadha, M., deSouza, L.A., Sterbing-D’Angelo, S.J., Moss, C.F., and Lumpkin, E.A. (2015). Somatosensory substrates of flight control in bats. Cell Rep 11, 851–858. 10.1016/j.celrep.2015.04.001.

Marshall, K.L., Clary, R.C., Baba, Y., Orlowsky, R.L., Gerling, G.J., and Lumpkin, E.A. (2016). Touch Receptors Undergo Rapid Remodeling in Healthy Skin. Cell Rep 17, 1719–1727. 10.1016/j.celrep.2016.10.034.

Marshall, K.L., Saade, D., Ghitani, N., Coombs, A.M., Szczot, M., Keller, J., Ogata, T., Daou, I., Stowers, L.T., Bonnemann, C.G., et al. (2020). PIEZO2 in sensory neurons and urothelial cells coordinates urination. Nature 588, 290–295. 10.1038/s41586-020-2830-7.

Meyers, J.R., MacDonald, R.B., Duggan, A., Lenzi, D., Standaert, D.G., Corwin, J.T., and Corey, D.P. (2003). Lighting up the senses: FM1-43 loading of sensory cells through nonselective ion channels. J Neurosci 23, 4054–4065.

Min, S., Chang, R.B., Prescott, S.L., Beeler, B., Joshi, N.R., Strochlic, D.E., and Liberles, S.D. (2019). Arterial Baroreceptors Sense Blood Pressure through Decorated Aortic Claws. Cell Rep 29, 2192–2201 e2193. 10.1016/j.celrep.2019.10.040.

Moayedi, Y., Duenas-Bianchi, L.F., and Lumpkin, E.A. (2018). Somatosensory innervation of the oral mucosa of adult and aging mice. Sci Rep 8, 9975. 10.1038/s41598-018-28195-2.

Murthy, S.E., Loud, M.C., Daou, I., Marshall, K.L., Schwaller, F., Kuhnemund, J., Francisco, A.G., Keenan, W.T., Dubin, A.E., Lewin, G.R., and Patapoutian, A. (2018). The mechanosensitive ion channel Piezo2 mediates sensitivity to mechanical pain in mice. Sci Transl Med 10. 10.1126/scitranslmed.aat9897.

Ranade, S.S., Woo, S.H., Dubin, A.E., Moshourab, R.A., Wetzel, C., Petrus, M., Mathur, J., Begay, V., Coste, B., Mainquist, J., et al. (2014). Piezo2 is the major transducer of mechanical forces for touch sensation in mice. Nature 516, 121–125. 10.1038/nature13980.

Shin, K.C., Park, H.J., Kim, J.G., Lee, I.H., Cho, H., Park, C., Sung, T.S., Koh, S.D., Park, S.W., and Bae, Y.M. (2019). The Piezo2 ion channel is mechanically activated by low-threshold positive pressure. Sci Rep-Uk 9. ARTN 6446 10.1038/s41598-019-42492-4.

Szallasi, A., and Blumberg, P.M. (1999). Vanilloid (Capsaicin) receptors and mechanisms. Pharmacol Rev 51, 159–212.

Truschel, S.T., Wang, E., Ruiz, W.G., Leung, S.M., Rojas, R., Lavelle, J., Zeidel, M., Stoffer, D., and Apodaca, G. (2002). Stretch-regulated exocytosis/endocytosis in bladder umbrella cells. Mol Biol Cell 13, 830–846. 10.1091/mbc.01-09-0435.

Witschi, R., Johansson, T., Morscher, G., Scheurer, L., Deschamps, J., and Zeilhofer, H.U. (2010). Hoxb8-Cre mice: A tool for brain-sparing conditional gene deletion. Genesis 48, 596–602. 10.1002/dvg.20656.

Woo, S.H., Lukacs, V., de Nooij, J.C., Zaytseva, D., Criddle, C.R., Francisco, A., Jessell, T.M., Wilkinson, K.A., and Patapoutian, A. (2015). Piezo2 is the principal mechanotransduction channel for proprioception. Nat Neurosci 18, 1756–1762. 10.1038/nn.4162.

Woo, S.H., Ranade, S., Weyer, A.D., Dubin, A.E., Baba, Y., Qiu, Z., Petrus, M., Miyamoto, T., Reddy, K., Lumpkin, E.A., et al. (2014). Piezo2 is required for Merkel-cell mechanotransduction. Nature 509, 622–626. 10.1038/nature13251.

Wu, J., Lewis, A.H., and Grandl, J. (2017). Touch, Tension, and Transduction -The Function and Regulation of Piezo Ion Channels. Trends Biochem Sci 42, 57–71. 10.1016/j.tibs.2016.09.004.

Zeng, W.Z., Marshall, K.L., Min, S., Daou, I., Chapleau, M.W., Abboud, F.M., Liberles, S.D., and Patapoutian, A. (2018). PIEZOs mediate neuronal sensing of blood pressure and the baroreceptor reflex. Science 362, 464–467. 10.1126/science.aau6324.

