## Supplemental Figures for "Lighting Up Mechanosensation: dyeing to see PIEZO2"

**Supplemental Figures, Villarino et al.**

SUPPLEMENTAL FIGURE 1

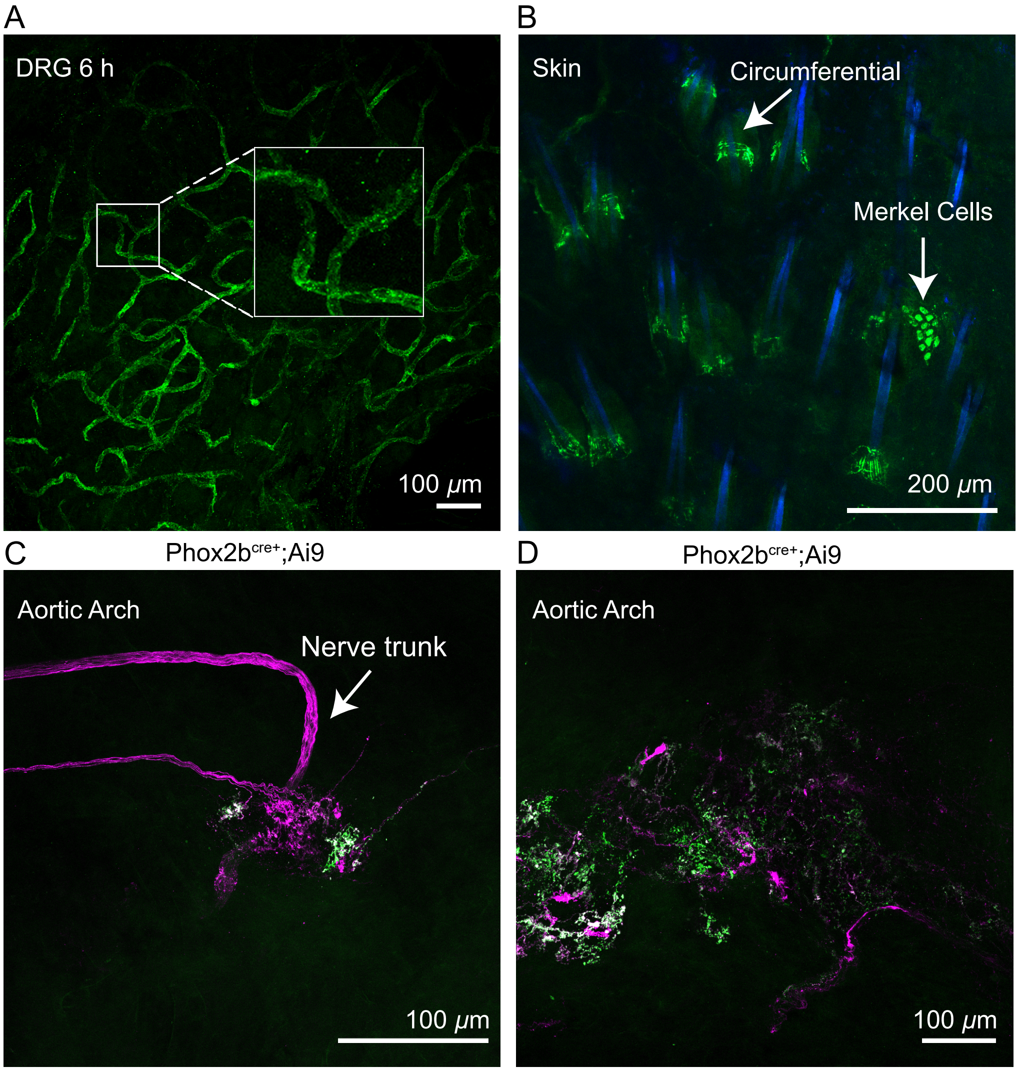


**Supplemental Figure 1: Patterns of FM 1-43 labeling**

(A) Zoomed-in inset of FM 1-43 labeling of vasculature in DRG at 6 h timepoint from Figure 3B. This labeling is no longer prominent after 12 hours.

(B) Representative z-stack image of 24 h FM 1-43 labeling of cutaneous neurons in whole-mount skin from wildtype mouse.

(C-D) Two z-stack images of examples of aorta baroreceptor endings from a Phox2b$^{Cre+}$;Ai9 mouse injected i.p. with FM 1-43 24 h prior to tissue collection. Tdtomato (magenta) and FM 1-43 (green) channels are displayed as a single merged image. Green endings are also tdTomato+, but some tdTomato+ endings did not label with FM 1-43.

SUPPLEMENTAL FIGURE 2

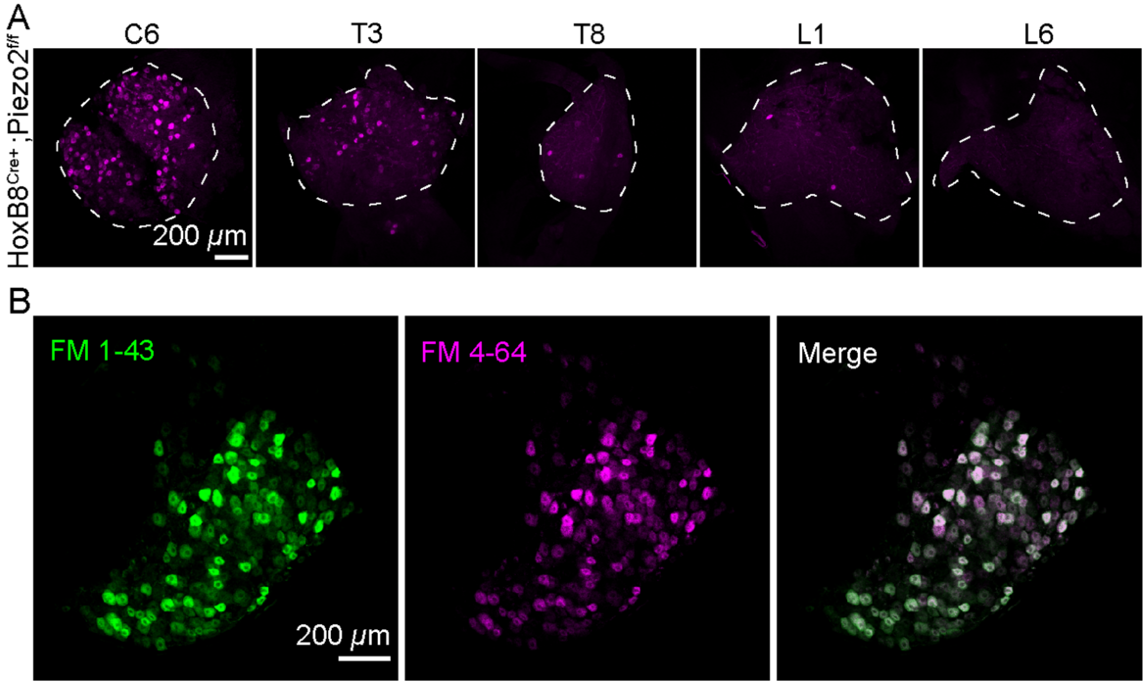


**Supplemental Figure 2: FM 4-64 sensory neuron labeling is consistent with that of FM 1-43**

(A) Representative z-stack images of whole-mount DRG from *HoxB8*$^{Cre+}$*;Piezo2*$^{f/f}$ mice that were injected with FM 4-64 24 h prior. The DRG level is indicated. Scale bar applies to all images.

(B) DRG from a wildtype mouse i.p. injected with both FM 1-43 (green) and FM 4-64 (magenta) 24 h prior to tissue harvest. Scale bar applies to all images.

SUPPLEMENTAL FIGURE 3

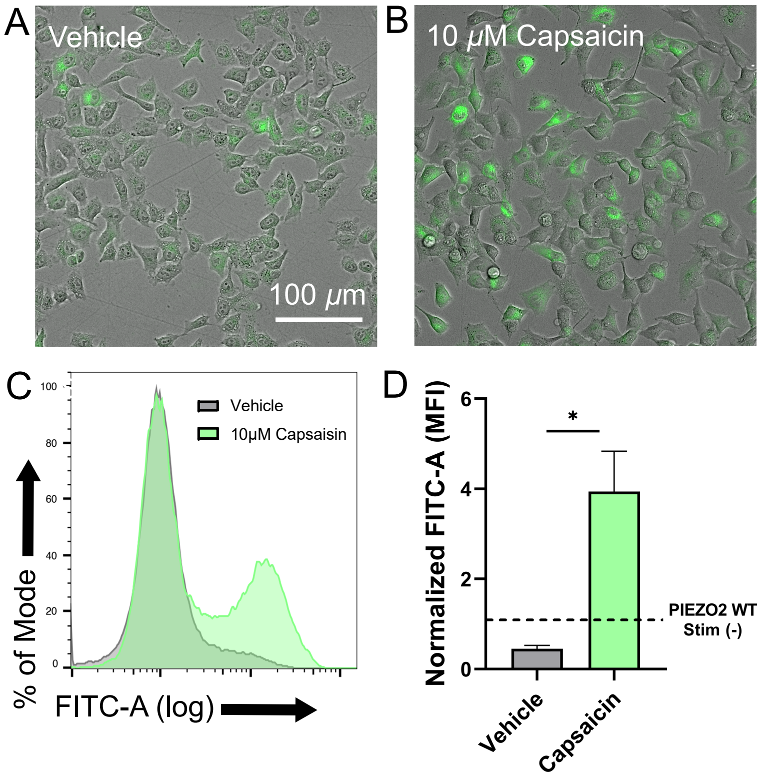


**Supplemental Figure 3: TRPV1 activity induces FM 1-43 labeling in cell culture**

(A) Representative image from HEKP1 KO cells transfected with TRPV1 treated with 10 μM FM 1-43 and vehicle control (EtOH).

(B) Representative image from HEKP1 KO cells transfected with TRPV1 treated with 10 μM FM 1-43 and 10 μM capsaicin.

(C) Flow cytometry histograms from FM 1-43 treated HEK P1KO cells. Vehicle (gray) and capsaicin (green) histograms are displayed as an overlay. X-axis is FITC-A (log) and represents FM 1-43 loading; Y-axis represents recorded events as a percentage of the mode.

(D) Flow cytometry quantification of each condition background subtracted and expressed relative to the levels observed in unstimulated HEK P1KO cells transfected with hPIEZO2 WT (shown as a dotted line).

SUPPLEMENTAL FIGURE 4

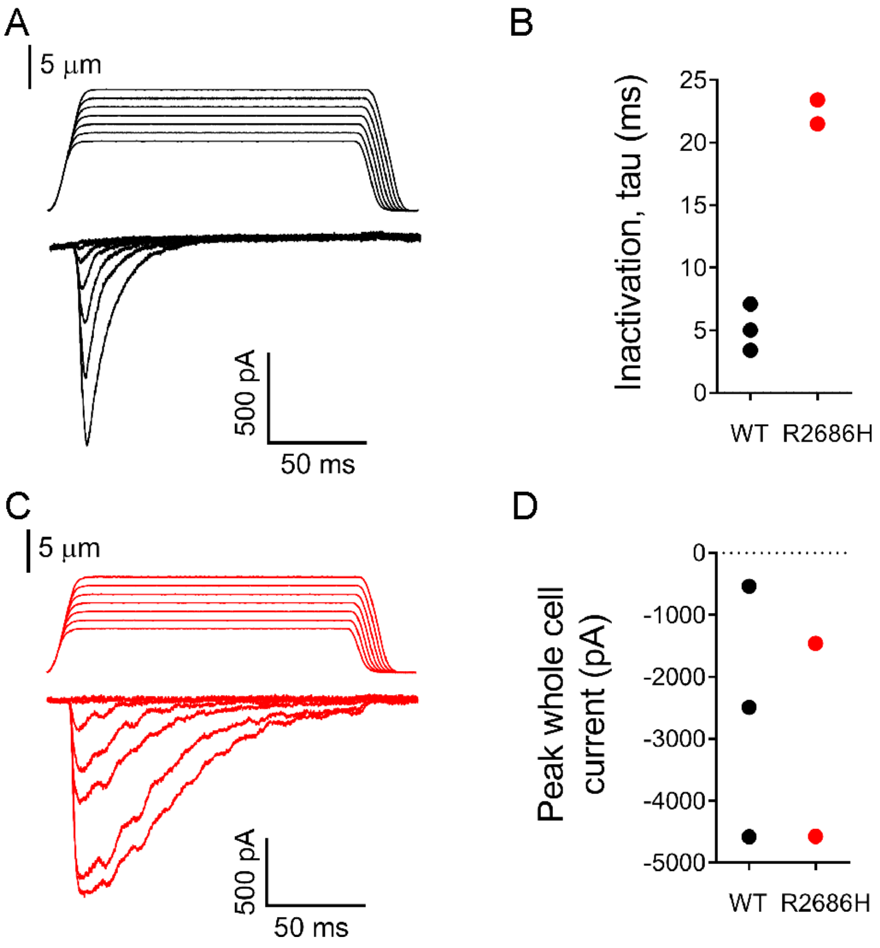


**Supplemental Figure 4:** ***PIEZO2-R2686H*** **exhibits strong gain-of-function inactivation phenotype**

(A) Poking induced whole cell currents for wildtype h*PIEZO2*. Stimulus traces (top) and current traces (bottom).
(B) Inactivation time constant tau for the indicated condition.
(C) Poking induced whole cell currents for h*PIEZO2* R2686H. Stimulus traces (top) and current traces (bottom).
(D) Peak whole cell currents from both conditions.

SUPPLEMENTAL FIGURE 5

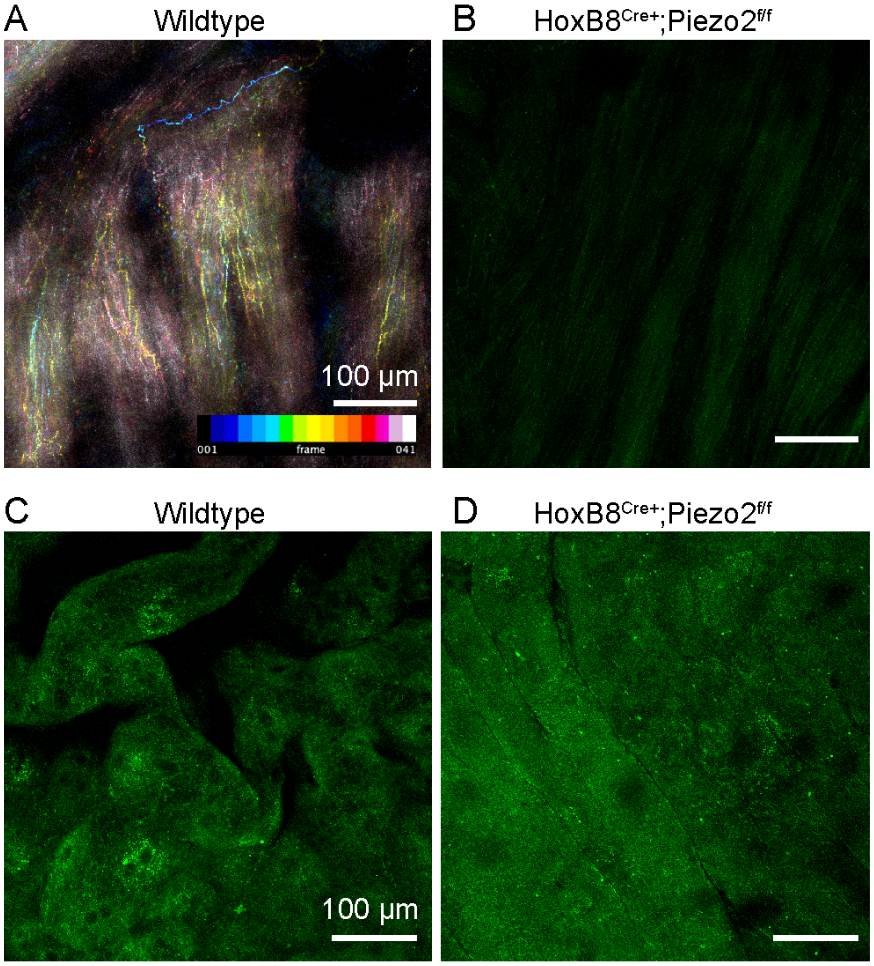


**Supplemental Figure 5: Bladder smooth muscle and urothelium labeling with FM 1-43**

(A) Representative z-stack of 41 images through the bladder of a wildtype mouse injected with FM 1-43 24 h prior. This image was taken from the muscle side. Stack was color bar coded by depth with blues indicating structures closer to the urothelium, and pink and white indicating structures within the muscle layer. A superficial neuronal process is visible in blue.

(B) Representative z-stack through the bladder of a *HoxB8*$^{Cre+}$*;Piezo2*$^{f/f}$ mouse injected with FM 1-43 24 h prior.

(C) Representative z-stack through the bladder, imaged from the urothelial side, in a wildtype mouse injected with FM 1-43 24 h prior.

(D) Representative z-stack through the bladder, imaged from the urothelial side, in a *HoxB8*$^{Cre+}$*;Piezo2*$^{f/f}$ mouse injected with FM 1-43 24 h prior.

SUPPLEMENTAL FIGURE 6

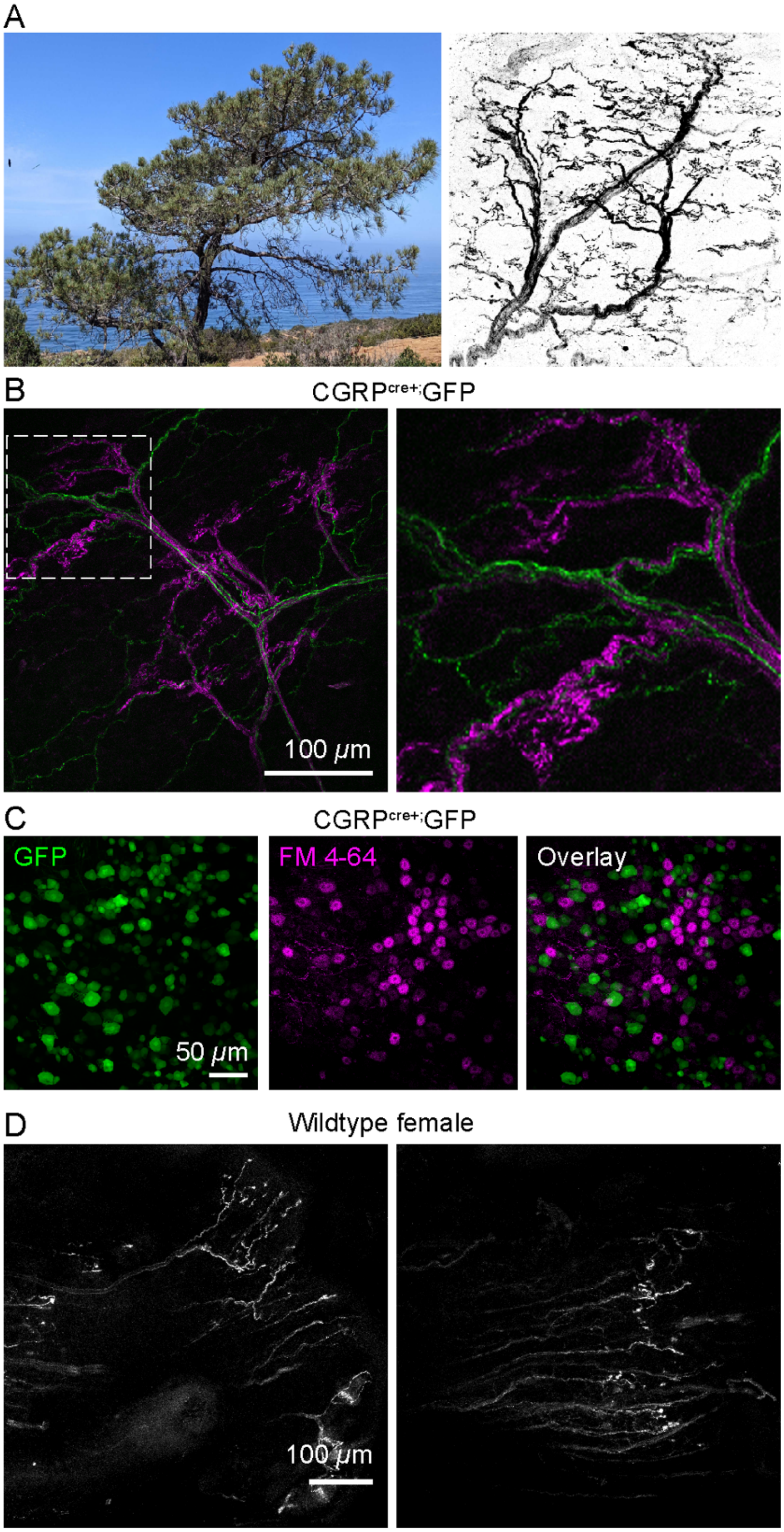

**Supplemental Figure 6: Characterization of Torrey Pines neurons and labeling in the distal urethra of female mice**

(A) Photograph of *Pinus torreyana* within Torrey Pines State Reserve (left) and representative inverted 20X image of Torrey Pines neurons from a wildtype mouse labeled with FM 1-43.

(B) Representative z-stack of a Torrey Pine neuron from a $CGRP^{cre+}$;GFP mouse, i.p. injected with FM 4-64 24 h prior. GFP (green) and FM 4-64 (magenta) channels are displayed as a single merged image. Image on right is zoomed-in from the boxed area on the left.

(C) Representative z-stack of a lumbar DRG from a $CGRP^{cre+}$;GFP mouse, i.p. injected with FM 4-64 24 h prior. GFP (green) and FM 4-64 (magenta) channels are displayed as a separate images. Image on right displays both channels as a single merged image.

(D) Two examples of FM 1-43 labeled ending structures at the distal end of the female urethra.
